# Evolutionary innovations in germline biology of placental mammals revealed by transcriptomics of first wave spermatogenesis in opossum

**DOI:** 10.1101/2023.06.17.545442

**Authors:** Kira L Marshall, Daniel J Stadtmauer, Jamie Maziarz, Günter P Wagner, Bluma J Lesch

## Abstract

Mammalian spermatogenesis is a deeply conserved developmental program that is essential for fitness. Paradoxically, spermatogenic development also allows rapid divergence in gene expression and is thought to be a source of evolutionary novelty and gene birth. How mammalian spermatogenic cells protect a conserved developmental program while enabling exceptionally rapid divergence in gene expression and function is unknown. Here, we comprehensively profile the spermatogenic gene expression program in grey short-tailed opossum (*Monodelphis domestica*, a model marsupial) and compare it to equivalent data from the mouse (*Mus musculus*, a model placental mammal) to discover contrasting forces underlying the unique evolutionary dynamics of gene expression during mammalian spermatogenesis. For the first time, we describe the timing of the ‘first wave’ of opossum spermatogenesis, and we combine bulk transcriptomic data from first-wave juvenile testes with single-cell transcriptomic data from adult testes to define conserved and divergent gene expression programs across the placental-marsupial split. We substantiate and extend our findings using genome-wide chromatin and multi-species transcriptome data and identify three classes of genes with different evolutionary trajectories: a deeply conserved central gene regulatory program governing spermatogenic progression; a separate class of spermatogenic genes exhibiting dynamic expression across placental mammals; and a third set of genes with evidence for directional selection in the placental mammal ancestor and constraint on expression levels within the placental mammalian lineage, representing placental innovations in germline gene expression and including biologically critical modules such as the DNA recombination and repair machinery.

## Introduction

New species-defining characteristics often emerge through divergence in spatial and temporal regulation of conserved developmental genes (Griffith et al. 2017; Lynch et al. 2008; Prabhakar et al. 2008; Prescott et al. 2015; Villar et al. 2015; King and Wilson 1975; Wray 2007; Carroll 2005). Discovery and quantification of gene regulatory divergence is therefore essential for understanding phenotypic evolution. However, comprehensive identification and quantification of gene regulatory differences requires careful matching of developmental stages, tissues, and cell types across species, creating practical challenges for robust discovery and validation of these changes in most biological systems.

Spermatogenesis — the process of male gamete development — is uniquely suited as a model system for addressing this and for understanding patterns of evolution of mammalian gene regulation. The tissue structure of the mammalian testis and developmental trajectory of spermatogenic differentiation are well conserved, but gene regulatory evolution is accelerated in the testis compared to other tissues (Good and Nachman 2005; Kopania et al. 2022; Swanson and Vacquier 2002; Turner et al. 2008; Brawand et al. 2011), facilitating identification of gene regulatory divergence within developmentally equivalent (homologous) cell types. Further, a reduced threshold for transcriptional activation during spermatogenesis leads to globally increased transcription and promotes expression of young or new transcripts (Kaessmann 2010; Xia et al. 2020; Soumillon et al. 2013), and changes in gene regulation that arise in the germline can be immediately passed to future generations (Assis and Bachtrog 2013; Findlay et al. 2009; Kaessmann 2010; Kondo et al. 2017; Marques et al. 2005; Xia et al. 2020).

Therefore, mammalian spermatogenesis provides an attractive framework to address a fundamental question in molecular evolution: how does evolutionary change in gene expression arise in the context of a deeply conserved developmental system? However, this developmental system has been underexploited for studying patterns of gene expression divergence, especially in mammalian evolution. Here, using histological, developmental, and transcriptomic characterization of adult and juvenile testes, we identify the timing of the ‘first wave’ progression of spermatogenesis in grey short-tailed opossum (*Monodelphis domestica*), a model marsupial that has been the source of many insights into genome and chromosome evolution (Gentles et al. 2007; Goodstadt et al. 2007; Mahony et al. 2007; Mikkelsen et al. 2007; Samollow 2008) and evolution of developmental systems (Chavan et al. 2021; Griffith et al. 2017; Lynch et al. 2008). The placental (eutherian) and marsupial (metatherian) lineages split from a common ancestor approximately 160 million years ago (Bininda-Emonds et al. 2007; Kumar et al. 2017; van Rheede et al. 2006). At the cellular level, placental and marsupial spermatogenesis is very similar, but several important molecular differences have been identified between placental and marsupial spermatogenesis; how these differences emerged and how they are maintained in the context of the deeply conserved spermatogenic program is unclear. To date only a single study, focusing primarily on somatic cell development, has carefully characterized male gonad development in *M. domestica* (Xie et al. 1996). Combining bulk transcriptomic data from the first wave and single cell transcriptomic data from adult testes, we identify robust gene expression differences between opossum and mouse, finding quantitative changes in expression of genes associated with known functional differences between marsupials and placental mammals and discovering previously undescribed differences in pathways important for reproduction. We apply sensitive, robust comparative methods to quantify divergence in expression of spermatogenesis-associated genes in opossum compared to mouse, test the generalizability of these changes across mammalian evolution, and classify genes according to patterns of conservation and divergence across the mammalian lineage.

We identify three classes of genes with different evolutionary trajectories: conserved genes, genes with dynamic expression patterns across species, and genes showing directional selection at the base of the placental mammalian lineage and constraint on expression in placental mammals. We mine this resource to uncover previously undescribed functional differences between placental mammal and marsupial germ lines, including regulation by WNT signaling and surveillance of reactive oxygen species. Our results establish a framework, model system, and resource dataset for studying the evolution of gene regulation and its contribution to evolution of reproduction and development.

## Results

### Conserved cell composition and transcriptional signatures in adult opossum testes

To define gene expression dynamics during spermatogenesis in opossum, we performed single cell RNA-sequencing (scRNA-seq, 10x Genomics) on dissociated whole testis samples from two adult opossums as well as two adult mice, after confirming that testis histology in the opossum visually conforms to the well-defined epithelial histology of mammalian spermatogenesis (**Figure 1A**) (Russell et al. 1990). We adapted a widely-used scRNA-seq analysis pipeline (Wolf et al. 2018) to account for challenges specific to the opossum data, including less mitochondrial gene expression and fewer unique gene identifiers (**Supplemental Figure 1A-B**). We tuned our pipeline to ensure retention of only high-quality reads from single cells and remove sperm head contamination (see methods) and combined replicates after confirming high similarity of clusters and quality metrics across replicates (**Supplemental Figure 1B-D**). Using established gene markers in mouse (Hermann et al. 2018; Shami et al. 2020; Wang et al. 2018; Murat et al. 2023; Green et al. 2018) that have orthologs in opossum, we identified clusters representing germ cells, somatic cells, and residual bodies/debris (**Figure 1B, Supplemental Figure 1E, 2, Supplemental Table 1**). Germ cells were re-clustered to identify fifteen and eighteen germ cell clusters for opossum and mouse, respectively (**Supplemental Figure 1E**), and clusters were assigned identities based on the same established marker genes used above (**Figure 1C, 1D** and **Supplemental Figure 3**). We labelled only populations we could assign with high confidence based on marker gene expression, leaving a transitional population in both species with no assigned identity that we removed from the dataset for all further analyses. Somatic cells were also re-clustered to identify eleven and nine somatic cell clusters for opossum and mouse, respectively (**Supplemental Figure 1E**), and clusters were assigned identities based on previously established marker genes (**Figure 1C** and **Supplemental Figure 4**) (Murat et al. 2023; Hermann et al. 2018; Shami et al. 2020). This final germ cell dataset of 2,990 cells in opossum and 9,112 cells in mouse and somatic dataset of 4,249 cells in opossum and 1,405 cells in mouse was used for downstream analysis.

**Figure 1:**
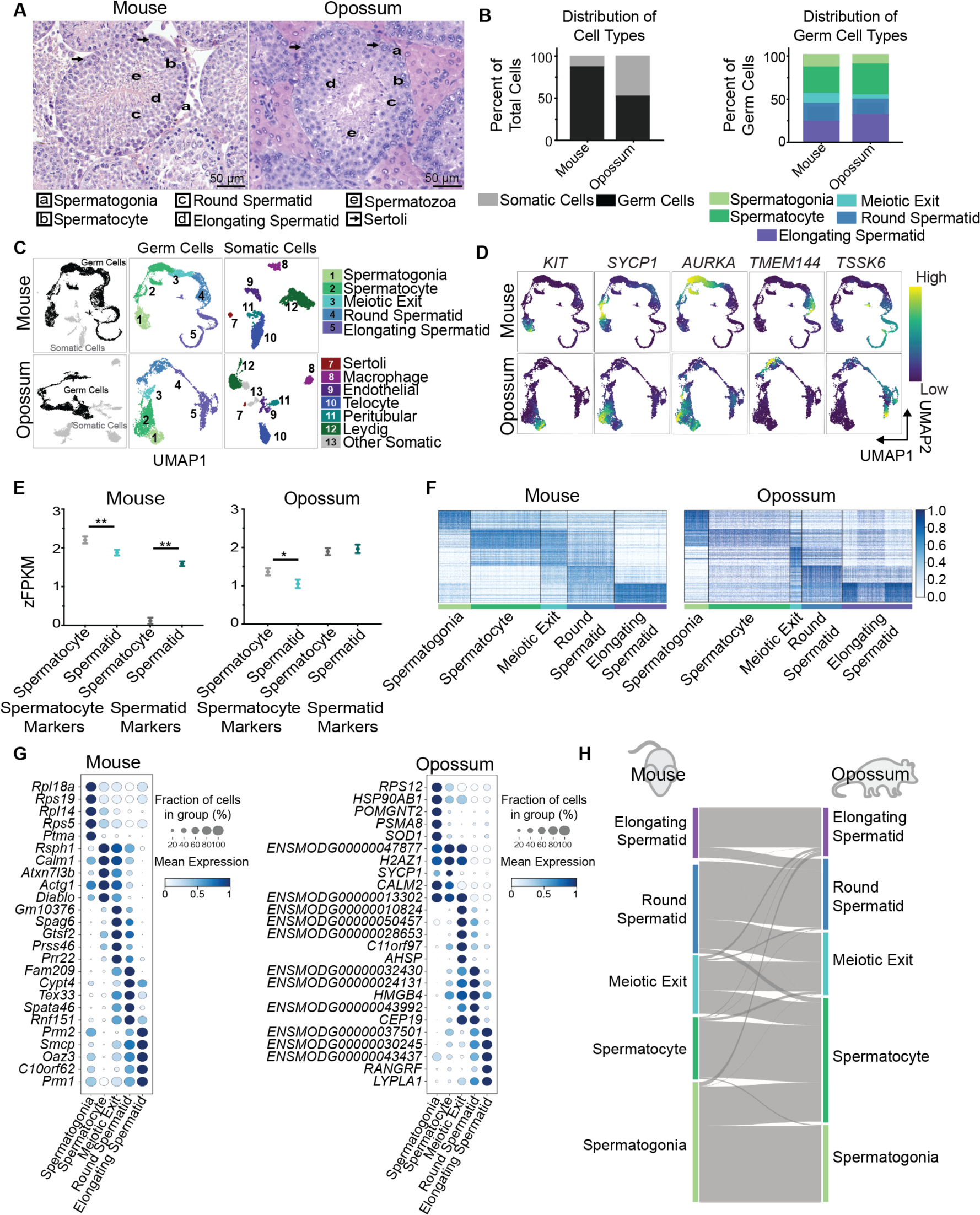
Single-cell analysis of adult mouse and opossum testis. A. Cross-sections of whole testis from adult mouse (left) or opossum (right) stained with hematoxylin and eosin. Representative cells of each type are indicated with letters (spermatogenic cells) or arrows (Sertoli cells). B. Percentage of cells identified as somatic vs. germ cells (left) and as specific spermatogenic cell types (right) in single cell clusters. C. Left: UMAP plots showing cell clusters (mouse: n = 10,953; opossum: n = 8,797) pseudocolored by somatic (grey) or germ (black) cell identity. Middle: Germ cell clusters pseudo-colored by spermatogenic cell type (mouse: n = 9,112; opossum: n =2,990). Right: somatic clusters pseudo-colored by cell type (mouse: n = 1,405; opossum: n = 4,249). D. Expression patterns of select cell markers for each spermatogenic cell type cluster for mouse and opossum projected on to the corresponding UMAP plot. Expression is shown as relative from low (purple) to high (yellow). For numerical scales of each gene, see Supplemental Figure 5. KIT: spermatogonia, SYCP1: spermatocytes, AURKA: spermatocytes, TMEM144: round spermatids, TSSK6: elongating spermatids. E. Distribution of zFPKM normalized expression values from sorted spermatocytes (grey) or spermatids (teal) for genes identified as markers of spermatocyte or round spermatid scRNA-seq clusters. Mean is plotted as a circle with error bars showing standard error of the mean (SEM). *p <0.05, **p <0.01 (Mann-Whitney U Test). F. Heatmap of expression for the top 100 marker genes for each germ cell cluster (mouse: 480 genes; opossum: 481 genes). Expression is scaled by gene from 0 to 1 (see Methods). G. Dot plot showing the top five marker genes for each cell type cluster. Size indicates the fraction of cells in each cluster expressing the gene; color indicates average expression of each gene per cluster. H. Sankey plot showing mapping of cell types between mouse (left) and opossum (right) scRNA-seq germ cell clusters. See Supplemental Table 1 for pairwise mapping scores.

Consistent with expectations and with our histology data, spermatogenesis in both species proceeds along a smooth trajectory from mitotically dividing spermatogonia to meiotic spermatocytes, followed by post-meiotic round spermatids, and finally elongating spermatids that are differentiating to sperm (**Figure 1C**). Our data revealed a higher percentage of somatic cells in opossum testes as compared to mouse, a surprising finding that we confirmed by histological examination (48.34% vs 12.83%; **Figure 1A, 1B**, see also **Figure 4B**). Germ cell clustering and overall cell type distribution was similar between opossum and mouse testes (**Figure 1C**). We also evaluated cell clustering and progression through spermatogenesis using Potential of Heat-diffusion for Affinity-based Trajectory Embedding (PHATE; (Moon et al. 2019)), an alternative visualization method designed to preserve local and global structure of the data. Marker gene expression in the PHATE projection shows similar progression between species and supports our assigned clusters (**Supplemental Figure 3E**).

We further validated cluster identities by comparing gene expression levels of the top 100 marker genes for the clusters identified as spermatocytes or round spermatids to normalized bulk RNA-seq data from sorted spermatocytes and round spermatids (Lesch et al. 2016). As expected, genes identified as markers of spermatocytes were expressed at significantly higher levels in sorted pachytene spermatocytes than round spermatids in both species (median 1.450 vs. 1.027, p =0.0423 and 2.166 vs. 2.002, p = 0.0110 for opossum and mouse, respectively; Mann-Whitney test; **Figure 1E**). The inverse pattern was also seen for round spermatids, though subtle and not statistically significant in opossum (median 1.797 vs. 1.925, p =0.5813 and 0.1599 vs. 1.544, p < 0.0001 in opossum and mouse, respectively; **Figure 1E**). Expression of the top 100 marker genes from each germ cell cluster across all germ cells (480 genes in mouse or 481 genes in opossum; **Figure 1F; Supplemental Table 1**) clearly separates the clusters. Clusters are also easily distinguished using only the top five markers (excluding mitochondrial genes) for each cell type, although many genes are expressed in multiple cell types (**Figure 1G**).

Direct comparison of cluster expression profiles between opossum and mouse using Self-Assembling Manifold mapping (SAMap; (Tarashansky et al. 2021)) showed strong quantitative agreement in expression profiles of single-cell clusters annotated as homologous cell types between species, confirming expectations that equivalent cell types in mouse and opossum are defined by highly conserved gene expression programs (**Figure 1H, Supplemental Table 1**). In each case, the highest-scoring mapping was between cell clusters manually identified as homologs. Between-species pairings were driven by biologically relevant gene pairs. For example, spermatogonia were linked by expression of *UCHL1*, spermatocytes by *SYCP3*, round spermatids by *KLF5*, and elongating spermatids were linked by multiple *TSSK* genes (**Supplemental Table 1**) (Hermann et al. 2018; Shami et al. 2020; Green et al. 2018). Most of the divergent edges between opossum and mouse clusters connect cell types immediately preceding or following in the temporal developmental trajectory, indicating shifts in the timing of expression and/or minor differences in cluster delimitation. These data support the accuracy of the assigned cluster identities and confirm broad similarities in the spermatogenic gene expression program between mouse and opossum.

### Timing of first-wave spermatogenesis in opossum

In mammals, neonatal testes contain only spermatogonial precursors, and a ‘first wave’ of spermatogenesis occurs during the first weeks of life as precursors differentiate, enter meiosis, and eventually produce sperm (Bellvé et al. 1977; McCarrey 1993). The first wave is a convenient system for studying spermatogenic development because new cell types are progressively added over time and gene expression signatures associated with each added cell type can be defined by comparing whole testes of progressively older animals. The transcriptomes of whole testes collected at specific developmental timepoints during the first wave of spermatogenesis can mimic the cell-type clustering in single cell analyses while allowing deeper and more robust sequencing coverage.

In the mouse, first-wave spermatogenic cells enter meiosis at 10 days postpartum (dpp), haploid round spermatids appear by 20 dpp, and mature sperm are present in the seminiferous tubule lumen by 35 dpp (Bellvé et al. 1977; McCarrey 1993). Because marsupials have a short internal gestation period and neonatal marsupials are altricial relative to neonatal placentals, it was unclear whether the opossum first wave would be initiated at birth, analogous to mice, or at a later time point when opossum joeys are developmentally equivalent to mouse neonates (Mate et al. 1994; Tyndale-Biscoe and Renfree 1987). A previous study established the presence of early meiotic cells in the testis at two months of age and mature sperm in the testis lumen at four months (Xie et al. 1996), but little additional detail is known.

To establish the timing of the first wave in opossum, we collected testes from juvenile male opossums at one month, two months, three months, and four months of age, as well as adults, and evaluated the seminiferous tubule morphology and range of cell types present in testes in tissue sections (**Figure 2A, B**). In seminiferous tubules of mammalian testis, spermatogenic precursors reside at the basal membrane and differentiate as they move toward the lumen; mature sperm produced at the end of differentiation are then released into the tubule lumen and carried to the epididymis. At one month, opossum seminiferous tubules showed a single layer of cells with no evidence of meiotic entry, inferred to be spermatogonia. By two months of age, early meiotic cells populate the tubules and pachytene spermatocytes were visible, however there was no evidence of cells progressing further than the first meiotic division. At three months, we observed the addition of small round cells with a single chromocenter, the characteristic appearance of post-meiotic round spermatids. By four months, tubules showed fully condensed spermatozoa in the lumen and were histologically comparable to adult testes. Additionally, by four months, mature spermatozoa were found throughout the epididymis, though male opossums are not considered sexually mature until five to six months (**Supplemental Figure 5A**) (Vandeberg 1989). The discrepancy between the presence of sperm and the timing of sexual maturity may be related to changes in seminiferous fluid, mating behavior, and/or maturation, as paired (mature) spermatozoa were present in the epididymis in adults but not at four months (**Supplemental Figure 5A**).

**Figure 2:**
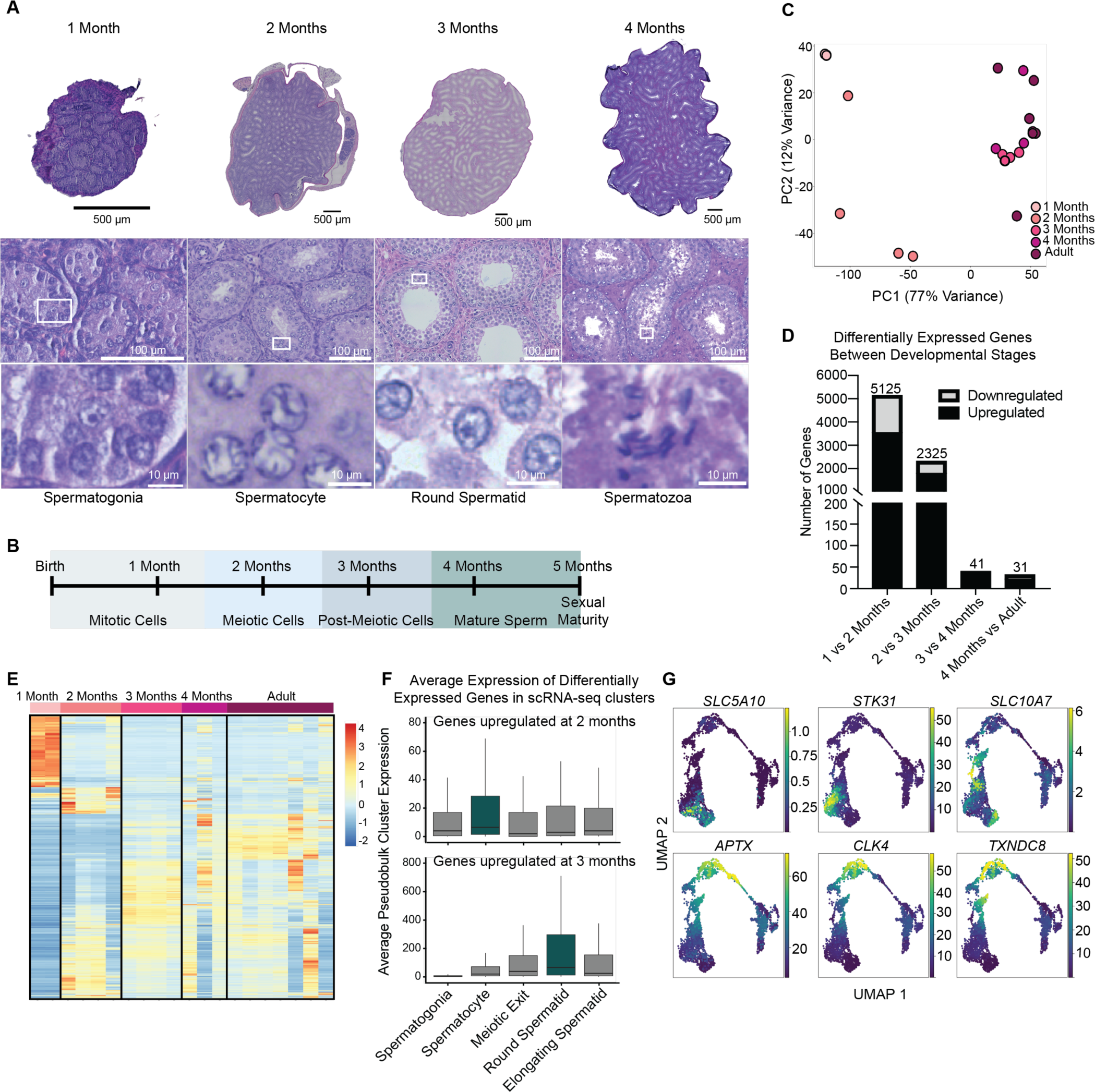
Characterization of developmental timing and differential gene expression during the first wave of spermatogenesis in opossum. A. Cross-sections of whole testis from opossum stained with hematoxylin and eosin at 1,2, 3, and 4 months. Images show whole testis (top), seminiferous tubules (middle), and examples of the most advanced spermatogenic cell type present at each age (bottom, regions from white boxes in middle images). The stage of the most advanced cells shown is indicated underneath the bottom row of images. B. Inferred timeline of progression through the first wave of spermatogenesis in opossum. C. Principal component analysis of bulk RNA-seq data from whole testis at each time point. Point color indicates opossum age: 1 month (n = 2), 2 months (n = 4), 3 months (n = 4), 4 months (n = 3), or Adult (n = 7). D. Pairwise comparisons of bulk RNA-seq data between consecutive developmental ages in opossum. Black indicates number of genes with higher expression in the older group and grey indicates number of genes with lower expression in the older group. Total number of differentially expressed genes for each comparison is shown above each bar. E. Heatmap displaying expression of all 7522 genes detected as differentially expressed in any comparison.^-^’ F. Average expression in scRNA-seq pseudobulk clusters of genes with increased expression from one to two months (top; 3283 genes) or two to three months (bottom; 1745 genes). Boxplots show interquartile range (IQR) with line at the median. Whiskers extend to 1,5*IQR. Teal box indicates the cluster representing the newly emerging cell type driving the increased expression during first-wave spermatogenesis. All differences are significant by Kruskal-Wallis and Dunn test (p<0.001) except Spermatogonia vs. Round Spermatid at 2 months and Meiotic Exit vs. Elongating Spermatid at 3 months. G. Select cell markers identified from bulk first-wave RNA-seq data as upregulated with the addition of meiotic cells (top) or post-meiotic cells (bottom) projected onto UMAP plots of scRNA-seq data.

### Transcriptional dynamics during opossum first-wave spermatogenesis reflect the progression of spermatogenic development

To define transcriptional dynamics during the first wave of opossum spermatogenesis, we collected bulk RNA-seq data from whole testes at each time point (n=2-7). Principal component analysis showed a distinct transcriptomic signature for both one-month and two-month timepoints, while three-month, four-month, and adult appear similar (**Figure 2C**). Hierarchical clustering of these datasets largely recapitulated these findings, with one-month and two-month time points clustering separately from the others (**Supplemental Figure 5B**).

We performed comparisons between each pair of consecutive ages to identify gene expression signatures associated with addition of each new spermatogenic stage in opossum. We filtered for genes with log2 fold change >2 between stages and an expression level of >50 reads across all samples to eliminate inflated fold change measurements due to low expression and extract high confidence differentially expressed genes (**Supplemental Figure 5C-D**). We identified 5,125 differentially expressed genes between one- and two-month old individuals, 2,325 DEGs between two and three months, 41 DEGs between three and four months, and 31 DEGs between four months and adult testes (**Figure 2D**, **Supplemental Table 2**). The very small differences observed in the last two time point comparisons are consistent with the expected cessation of transcription after the round spermatid stage (Kierszenbaum and Tres 1978) (which is already present at three months), with the histologic similarity between four month and adult testes, and with the similarity between datasets observed by PCA and hierarchical clustering (**Figure 2C, Supplemental Figure 5B**). As expected, the majority of differentially expressed genes were upregulated in the later age group as compared to the earlier for each comparison, as new cell types are added at later time points but none are lost. Gene Ontology analysis compared to a background of all genes expressed in either age group (**Supplemental Figure 5E-F**) confirmed that differentially regulated genes had functions consistent with the stage of spermatogenic cell added at the later time point. For example, comparison between one- and two-month testes highlighted transposition and microtubule-related pathways related to the onset of meiosis and transposon activation (Branciforte and Martin 1994; Kallio et al. 1998), while genes upregulated between two and three months included the categories “spermatid differentiation” and “spermatid development” (**Supplemental Figure 5E-F**). Consistent with these results, distinct gene clusters associated with each stage of gametogenesis were evident in the set of 6,623 genes differentially expressed in each pairwise comparison (**Figure 2E**). As expected, distinct transcriptional states were most evident between the early pairs of time points, when most transcriptional differences occur.

### Transcriptional dynamics during opossum first-wave spermatogenesis recapitulate gene expression patterns observed by scRNA-seq

To confirm that our scRNA-seq and bulk first-wave RNA-seq data yield comparable results, we compared the differentially expressed genes from bulk data at developmental timepoints where meiotic or post-meiotic cells were added to the average gene expression per scRNA-seq cluster in our adult single cell analysis (**Figure 2F, Supplemental Figure 5G-H**). We calculated pseudobulk average gene expression values for each scRNA-seq cluster and evaluated expression of genes upregulated at two months or three months in our bulk first-wave RNA-seq data. Genes upregulated at two months with the transition to meiotic spermatocytes showed highest expression in the single cell cluster corresponding to spermatocytes (p < 0.001 for all pairwise comparisons; **Figure 2F**). Genes upregulated at three months with the addition of round spermatids showed highest expression in the round spermatid cluster (p < 0.001 for all pairwise comparisons; **Figure 2F**). Similar pseudobulk analysis using finer division of our “spermatocyte”, “round spermatid”, and “elongating spermatid” clusters into two clusters each, representing early and late stage within each cell type, recapitulated these results, with genes with increased expression at the two-month and three-month timepoints showed highest average expression in the “spermatocyte” and “late round spermatid” clusters, respectively (**Supplemental Figure 5G, 3**). A separate comparison of the top 100 marker genes for each of the finer cell clusters to the sets of genes upregulated at the two- and three-month time points further supports a good correspondence between cell type-specific gene expression profiles detected in the scRNA-seq and bulk first-wave datasets (**Supplemental Figure 5H**, **Supplemental Table 2**).

Select genes identified as differentially expressed during the first wave showed expression at corresponding stages in UMAP plots of our scRNA-seq data (**Figure 2G**). Together, these data indicate that cell type-specific expression inferred from bulk first-wave and single cell adult testis transcriptomic data are comparable.

Differences between the datasets, especially at the meiotic stage, may represent contributions of developmental changes in gene expression in spermatogonia or somatic cells to the bulk data. To assess the somatic cell contribution, we performed a similar comparison by identifying the number of total somatic cell cluster markers in our scRNA-seq analysis that were differentially expressed at the two developmental transitions highlighted above and found that 106 (17.85%) and 74 (12.46%) of the 594 marker genes were upregulated from one to two months or two to three months, respectively, isolating a contribution from testis somatic cells to postnatal bulk gene expression changes (**Supplemental Table 2**).

### Quantitative cross-species comparison of the spermatogenic transcriptome

While single cell data are useful for comparisons of cell type markers and developmental trajectories between species, quantitative comparisons of expression across species can be more reliably performed using bulk RNA-seq data, especially for lowly expressed genes, due to the better coverage depth per gene (Squair et al. 2021; Zhang et al. 2020; Zheng et al. 2017; Kim et al. 2020). Since we found that cell types identified by scRNA-seq are comparable between mouse and opossum and confirmed that our first-wave bulk RNA-seq data accurately reflects these cell types, we next leveraged our first-wave bulk RNA-seq dataset to discover quantitative species differences in gene expression between opossum and mouse during spermatogenesis (**Figure 3A**).

**Figure 3:**
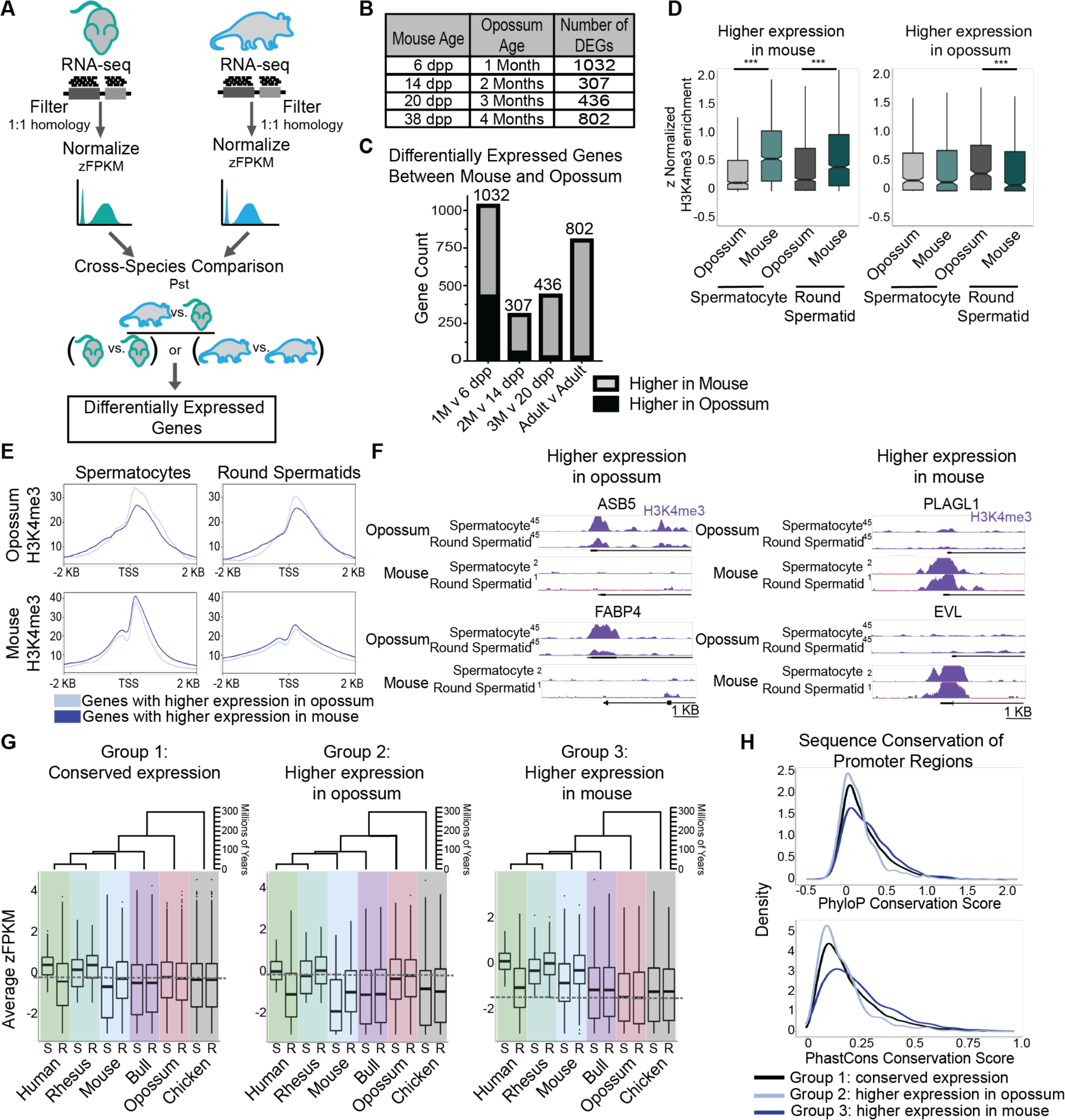
Differentially expressed genes between mouse and opossum reveal evolutionary innovations in placental mammalian gene expression. A. Strategy for comparing RNA-seq data across species. B-C. Summary of matched mouse and opossum developmental ages and numbers of differentially expressed genes in pairwise comparisons at each stage. In C, the total number of differentially expressed genes for each pairwise comparison is shown above each bar. D. Distribution of z-normalized H3K4me3 enrichment in sorted spermatocytes or round spermatids from mouse or opossum for genes found to be differentially expressed between mouse and opossum. Boxplots show interquartile range (IQR) with notch at the median. Whiskers extend to 1.5¶QR. ***p <0.0001 (Mann-Whitney U Test). E. Metagene plots of H3K4me3 signal in opossum or mouse sorted spermatocytes or round spermatids at promoter regions of genes with higher expression in the indicated species. F. Genome browser tracks showing H3K4me3 ChIP-seq enrichment in sorted spermatocytes or round spermatids from opossum or mouse at genes found to be differentially expressed between species. PLAGL1: Chr2:422,126,000-422,132,000 (opossum) or Chr10:13,088,000-13,094,000 (mouse); ASB5: Chr5:91,620,000-91,626,000 (opossum) or Chr8:54,548,000-54,554,000 (mouse); FABP4: Chr3:151,895,000-151,911,000 (opossum) or Chr3:10,205,000-10,211,000 (mouse); EVL: Chr1:318,927,000-318,133,000 (opossum) or Chr12:108,552,000-108,558,000 (mouse). Scale bar, 1 kilobase. G. Distribution of zFPKM values of genes showing conserved expression (Group 1 ; left), higher expression in opossum (Group 2; middle), or higher expression in mouse (Group 3; right) from sorted spermatocytes (S) or round spermatids (R) across four placental, one marsupial (opossum), and one other amniote species (chicken) as an outgroup. Boxplots show median and IQR; whiskers extend to 1 5*IQR. The relevant phylogenetic tree is shown above each graph. Dashed grey line indicates median of opossum round spermatid for reference. H. Distributions of PhyloP (top) and PhastCons (bottom) conservation score for promoter regions (± 1 kb from transcription start site) of genes showing conserved expression (Group 1 ; black), higher expression in opossum (Group 2; light blue), or higher expression in mouse (Group 3; dark blue) across 30 placental mammals, 1 marsupial, and 4 other vertebrates. Negative and positive PhyloP scores indicate faster or slow than expected evolution of DNA sequence, respectively, and a higher PhastCons conservation score indicates stronger sequence conservation across the 35 species

For comparison to mouse, we used a published high-time-resolution first-wave bulk RNA-seq dataset (Margolin et al. 2014). We limited our analysis to transcripts of genes with one-to-one orthology between mouse and opossum (13,521 genes; **Supplemental Table 3**) and normalized the data within each individual to obtain zFPKM values (Hart et al. 2013). zFPKM normalization is a robust method for cross-species RNA-seq analysis that is well suited to datasets representing varied complexity and time-series information (Hart et al. 2013; Uebbing 2018).

To evaluate differential gene expression between species, we applied Pst, a metric that compares sample variation among individuals of the same group to that of all individuals between groups (Antoniazza et al. 2010; Uebbing et al. 2016). We calculated Pst values for each gene at matched time points across species and obtained p-values by bootstrapping (see methods). To isolate high-confidence differentially expressed genes, we filtered based on both Benjamini-Hochberg adjusted p-value and between-species variation. To establish a filtering threshold for between-species variation, we examined control genes with known biological roles and expression patterns that have a high probability of being similar or different between species; this filter allows us to exclude genes where low within-species variation results in artificially inflated Pst values (**Supplemental Figure 6A-B**). The housekeeping genes eukaryotic elongation factor-2 (*EEF2*) and β-actin (*ACTB*) are expected to have consistent expression patterns across species, while *PRDM9* and *ZCWPW1* are known to be differentially expressed (Cavassim et al. 2022). PRDM9 is a histone lysine methyltransferase with a critical role in meiotic recombination in mouse, but it is functionally inactive in opossum due to a deletion of the DNA binding domain (Baker et al. 2017; Oliver et al. 2009). ZCWPW1 is a PRDM9 cofactor. *PRDM9* and *ZCWPW1* are highly expressed in mouse meiotic cells, but nonfunctional and lowly expressed during opossum spermatogenesis. Using these genes as negative and positive controls, we set a stringent between-species variation cutoff of greater than 3.5 in order to maximize specificity and minimize false-positive hits. We called differentially expressed genes using the filters described above and with a significance threshold of p<0.05 after Benjamini-Hochberg correction.

Next, to ensure that this normalization step did not create systematic artifacts, we examined the dynamics of genes with known, well-conserved functions in pre-meiotic, meiotic, and post-meiotic stages. These developmentally dynamic genes displayed expected stage-specific changes in expression through development, and dynamics were similar in both species. (**Supplemental Figure 6C-E**).

### Quantification of gene expression differences during mouse and opossum spermatogenesis

We identified 1032, 307, and 436 genes as differentially expressed between mouse and opossum at time points corresponding to emergence of pre-meiotic precursors (spermatogonia), early meiotic cells (spermatocytes), and post-meiotic haploid cells (spermatids), respectively (**Figure 3B**-**C, Supplemental Table 3**). While representing a minority of the genes identified as differentially expressed, 472 genes showed higher expression in opossum compared to mouse throughout the first wave (**Figure 3C, Supplemental Table 3**). Seventy genes were differentially expressed between species at all developmental timepoints examined (**Supplemental Table 3**), while 132 genes were shared between all comparisons starting at the time point corresponding to emergence of meiotic cells (two-month opossum versus 14 dpp mouse) (**Supplemental Table 3**). The lower number of differentially expressed genes shared across all time points relative to individual time points may be a dilution effect as the total number of cell types increases with age and decreases sensitivity for detecting subtle quantitative differences. Analysis of first-wave data therefore facilitates identification of cell type-specific differences that are masked in the adult whole testis while benefiting from the greater sequencing depth of bulk compared to single cell data. Examples of genes we detected as differentially expressed at all time points include spermatogonial stem cell homeostasis regulators *NEDD4* and *LIN28A* (Chakraborty et al. 2014; Wang et al. 2020; Zheng et al. 2009; Zhou et al. 2017), both of which are more highly expressed in mouse relative to opossum (**Supplemental Figure 6F**). Their appearance as differentially expressed genes in all comparisons is consistent with the presence of spermatogonia across each time point.

### Species-specific gene expression patterns reflect underlying species differences in chromatin state

To validate if the observed differences in gene expression and determine if they reflect species differences in underlying regulatory machinery, we used our previously published chromatin immunoprecipitation and sequencing (ChIP-seq) data in sorted germ cells to compare enrichment of the activating mark trimethylation of lysine 4 on histone 3 (H3K4me3) in promoter regions of genes showing differential expression (Lesch et al. 2016). As predicted, for genes identified as more highly expressed in mouse, H3K4me3 enrichment was significantly higher in germ cells of mouse compared to opossum (median 0.1331 vs. 0.5552, p < 0.0001 and 0.1894 vs. 0.4119, p < 0.0001 for spermatocytes and spermatids, respectively; Mann-Whitney test; **Figure 3D-E**). The reverse was also true for genes more highly expressed in opossum, though not statistically significant in sorted spermatocytes (median 0.1636 vs. 0.1322, p = 0.84 and 0.2836 vs. 0.0820, p < 0.0001 for spermatocytes and spermatids, respectively; **Figure 3D-E**). Visualization of H3K4me3 enrichment at individual differentially expressed genes on the genome browser further confirmed robust species-specific differences in chromatin state (**Figure 3F**). These results reinforce the validity of the identified expression differences and suggest that these differences reflect underlying divergence of regulatory state in the mammalian germ line.

### Opossum-mouse divergence identifies evolutionary innovations in placental mammalian gene expression

We next asked if the identified differences in expression between opossum and mouse reflect more general patterns across the mammalian lineage. For genes identified as having conserved or divergent expression in mouse and opossum, we compared zFPKM-normalized gene expression levels in sorted spermatocytes and round spermatids from five mammalian (human, rhesus macaque, mouse, bull, opossum) and one amniote (chicken) species spanning approximately 300 million years of divergence (**Figure 3G**) (Lesch et al. 2016; Kumar et al. 2017; Bininda-Emonds et al. 2007; Brocklehurst et al. 2022). As predicted, genes with conserved expression between mouse and opossum (11,701) showed global conservation in expression level in spermatogenic cells across amniotes (**Figure 3G**). Genes with higher expression in opossum compared to mouse (472 genes) demonstrate varying expression patterns across the Boreoeutherian (scrotum-carrying; includes bull, mouse, rhesus macaque, and human) lineage. This pattern is consistent with dynamic adaptive gene expression levels with increasing variability across all therian mammals, rather than marsupial-specific divergence. In contrast, genes with higher expression in mouse compared to opossum (1,350 genes) were consistently more highly expressed in other placental mammal species, suggesting directional selection for increased expression of these genes during spermatogenesis in the placental mammalian ancestor and constraint on expression levels within the placental mammalian lineage. To provide further support for these patterns, we evaluated sequence conservation in regulatory regions for each class of genes by comparing mean PhyloP and PhastCons conservation scores in promoter regions (transcription start site ± 1kb) using alignments from 30 placental, 1 marsupial, and 4 other vertebrate species (**Figure 3H**) (Blanchette et al. 2004; Siepel et al. 2005). Promoters of genes with unstable expression during placental evolution had reduced sequence conservation and higher than expected rates of base pair evolution compared to the promoters of conserved and placental-elevated groups (p < 0.001, Kruskal-Wallis and Dunn test for multiple comparisons). Conversely, consistent with a history of directional selection at the base of the placental mammalian lineage and stabilizing selection thereafter, promoters of genes with elevated expression in mouse compared to opossum have higher sequence conservation in the placental-dominated alignment (p < 0.001, Kruskal-Wallis and Dunn test for multiple comparisons). These data highlight three evolutionarily distinct spermatogenic gene expression programs: a deeply conserved central gene regulatory program governing spermatogenic progression (Group 1), a set of genes showing dynamic expression levels across mammalian spermatogenesis (Group 2), and a third group of genes representing placental innovations in testis gene expression that have undergone selection for higher expression in the placental lineage (Group 3).

### Validation of differentially expressed genes in vivo

To better understand the biological significance of species differences in expression, we selected a subset of genes to further investigate *in vivo*. We first performed reverse transcription followed by quantitative PCR (RT-qPCR) in adult testes for three genes we identified as differentially expressed at multiple developmental timepoints and one control gene that showed no difference in expression between opossum and mouse whole testis across ages to confirm our differential expression calls in an independent set of samples (**Figure 4A**). The increased sensitivity of RT-qPCR also avoids the dropout problem inherent in high-throughput RNA sequencing. There was good correspondence between RT-qPCR results and predictions based on comparative transcriptomic analysis. Two additional genes called as differentially expressed did not meet significance by qPCR but showed a strong trend toward differential expression in the expected direction (**Supplemental Figure 6G**). We then selected two differentially expressed genes, *PIWIL2* (higher in mouse) and *HEXB* (higher in opossum), and visualized their tissue localization in mouse and opossum testis sections using single-molecule in-situ hybridization (sm-ISH; (Wang et al. 2012)). As predicted based on its known function in regulation of the piRNA pathway, *PIWIL2* transcripts were concentrated in meiotic cells in both species. *HEXB* was broadly expressed in most germ cells in opossum but was not seen in opossum interstitial cells and was virtually absent in all mouse cells (**Figure 4B**). While comparison across species using RNA FISH is not absolutely quantitative due to the unavoidable use of different probes, we observed a greater concentration of chromogenic stain for *PIWIL2* in mouse testes compared to opossum, and a greater concentration of *HEXB* in opossum compared to the mouse (**Figure 4B**) consistent with our evidence from RNA-seq and qPCR analyses. Together, high time-resolution qPCR and sm-ISH provided additional validation of identified species differences in gene expression.

**Figure 4:**
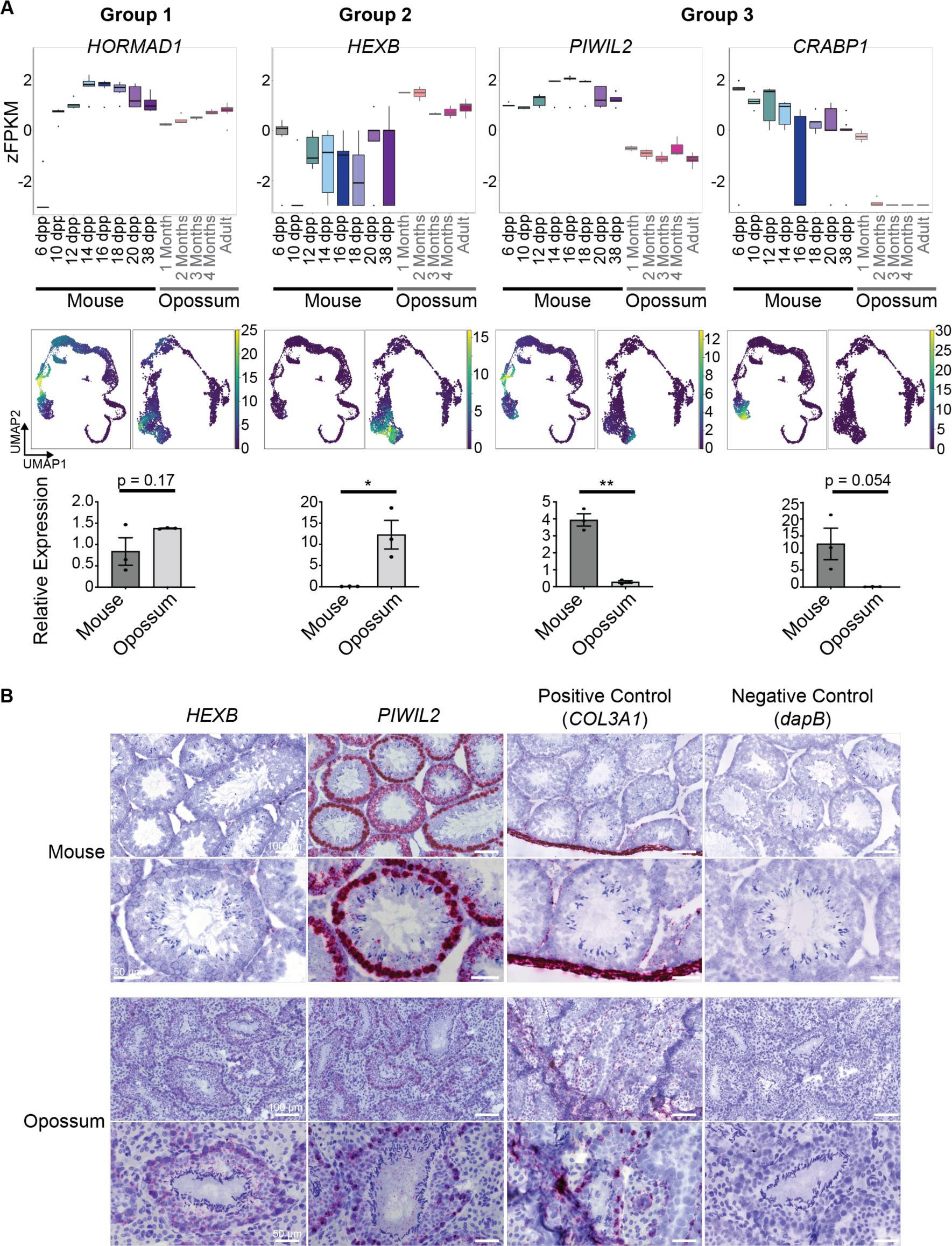
Confirmation of species differences in expression in testis tissue in vivo. A. Top: Distribution of zFPKM values at each developmental stage for four representative genes falling in different groups. HORMAD1, no difference (Group 1); HEXB, higher in opossum (Group 2); PIWIL2 and CRABP1, higher in mouse (Group 3). Mouse: 6 days post-partum (dpp; n = 6), 10 dpp (n = 5), 12 dpp (n = 6), 14 dpp (n = 6), 16 dpp (n = 13), 18 dpp (n = 6), 20 dpp (n = 5), or adult (i.e., 38 dpp; n = 8). Opossum: one month (n = 2), two months (n = 4), three months (n = 4), four months (n = 3), adult (n = 7). Middle: UMAP projections from scRNA-seq of the same four representative genes. Bottom: Relative expression of each gene in adult whole testis determined by qRT-PCR. Error bars = SEM, *p < 0.05, **p < 0.01 (unpaired t-test). B. mRNA expression by RNAScope (pink) with hematoxylin counterstain (purple) in testis of mouse or opossum for a representative Group 2 gene (HEXB) and a representative Group 3 gene (PIWIL2). Positive control is collagen gene COL3A1 (expressed in telocytes and fibroblasts, and accumulated in the tunica albuginea in both mouse and opossum) and negative control is bacterial gene dapB. Top row of images for each species shows multiple and bottom row shows single seminiferous tubules at higher magnification.

### Differences in gene expression levels at specific developmental stages reveal divergently regulated biological mechanisms in spermatogenesis

To further explore the biological importance of divergence in expression, we turned to the ontogenetic information inherent in our first-wave data. At different developmental time points, systematic differences between opossum and mouse gene expression correspond to functions known to be important at the relevant developmental stage and highlight potential functional differences between species. For example, at the time point when only pre-meiotic precursors (spermatogonia) are present in the testes (one-month opossum (n = 2) vs. 6 dpp mouse (n = 6)), genes belonging to the GO term “Wnt signaling” were enriched among this gene set (adjusted p=0.017; **Figure 5A**), consisting of eighteen differentially expressed genes that participate in the WNT signaling pathway, including *FZD5*, *DVL1*, and *DVL3* (**Figure 5A**; **Supplemental Table 4**). WNT signaling has been implicated in primordial germ cell specification, spermatogonial stem cell proliferation, and retention of stem cell state in mouse and human (Chassot et al. 2017; Golestaneh et al. 2009; Ohinata et al. 2009; Chawengsaksophak et al. 2012). Virtually nothing is currently known about how germline stem and progenitor cells differ between placentals and marsupials, and our results imply that altered WNT signaling may underlie divergence in regulatory pathways controlling spermatogonial stem cell function between the placental or marsupial lineages.

**Figure 5:**
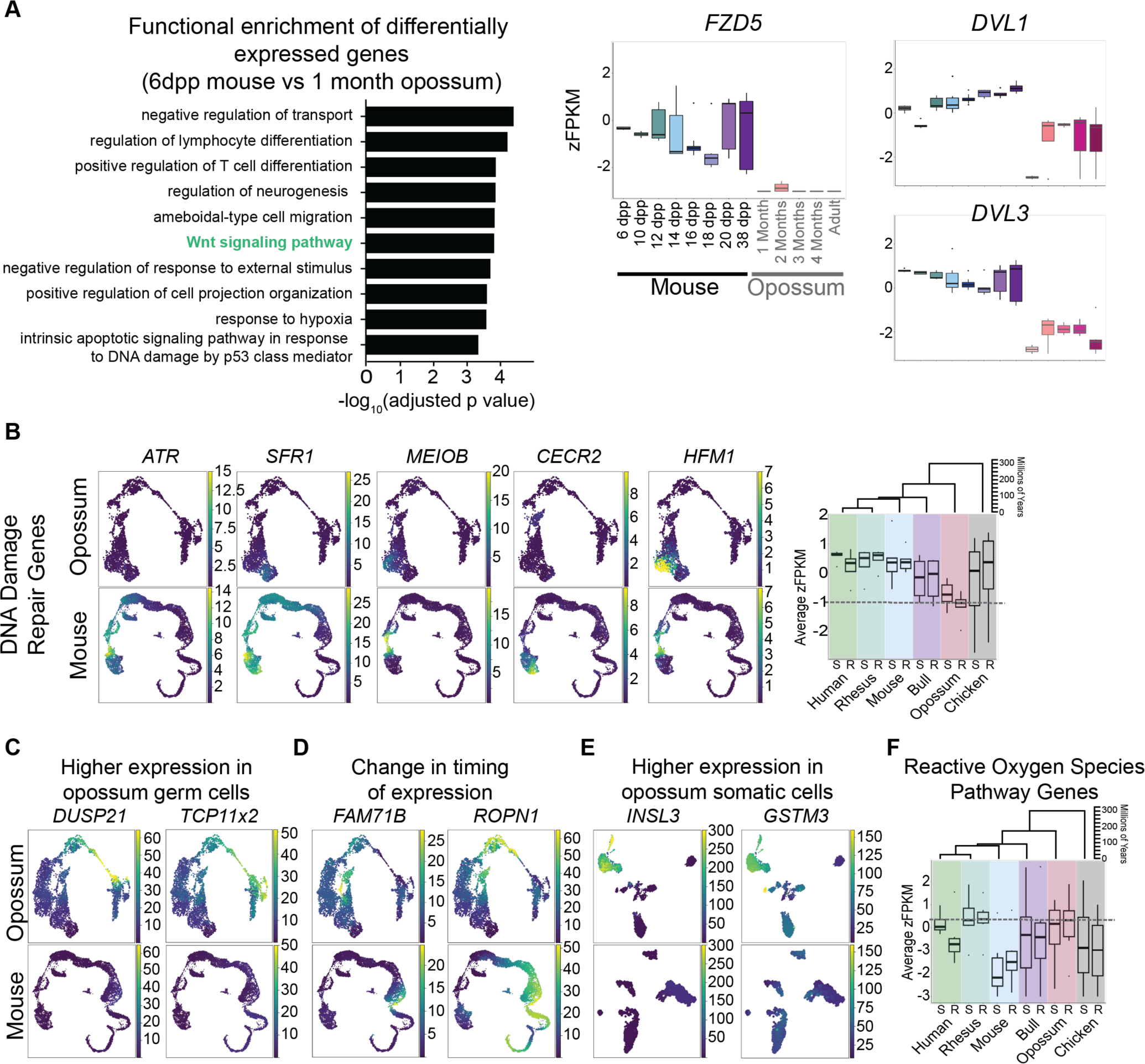
Biological roles for evolutionary innovations in gene expression during spermatogenesis. A. Left: Gene Ontology categories enriched among differentially expressed genes between one-month old opossum and six-day old mouse. Right: Distribution of zFPKM values in mouse and opossum first-wave datasets for three WNT signaling genes with higher expression in mouse compared to opossum throughout spermatogenesis. Boxes show the distribution of values across all replicates (n=2 to 13) at each time point. Dpp= days post-partum (mouse). B. Expression patterns in scRNA-seq of selected genes with higher expression in mouse (bottom) compared to opossum (top) representing components of DNA damage repair and homologous recombination pathways. Right: Distribution of zFPKM values of the same five DNA damage repair genes in sorted spermatocytes (S) or round spermatids (R). Boxplots display median and IQR; whiskers show 1,5*IQR. The relevant phylogenetic tree is shown above the graph. C-E. Expression patterns in scRNA-seq data for selected genes with higher expression in opossum compared to mouse. C. Two genes demonstrating higher expression in germ cells of opossum relative to mouse. D. Two genes demonstrating alternative timing of expression during opossum vs. mouse spermatogenesis. E. Two genes demonstrating higher expression in somatic cells of opossum relative to mouse. F. Distribution of zFPKM values of six genes identified as having higher expression in opossum with a role in reactive oxygen species processing (EPHX2, FOS, MSRA, NQO1, PTGS2, and GPX2) in sorted spermatocytes (S) or round spermatids (R). Boxplots display median and IQR; whiskers show 1 5*IQR. The relevant phylogenetic tree is shown above the graph.

In a second example, at time points following the development of meiotic cells (two-month-old opossum (n = 4) and 14 dpp mouse (n = 6)), many of the differentially expressed genes, including *ATR, SFR1, MEIOB, CECR2*, and *HFM1* (**Figure 5B**, **Supplemental Figure 6F, Supplemental Table 3**), are important for meiotic recombination and DNA damage repair (Akamatsu and Jasin 2010; Guiraldelli et al. 2013; Lee et al. 2012; Luo et al. 2013; Souquet et al. 2013; Widger et al. 2018), consistent with higher rates of meiotic recombination reported in placental compared to marsupial mammals (Mikkelsen et al. 2007; Samollow 2008; Samollow et al. 2007). The differentially expressed genes involved in recombination and DNA damage repair show a consistent gain in expression in placental mammal species compared to opossum (Group 3), suggesting that an increase in these pathways during spermatogenesis is a synapomorphy of placental mammals (**Figure 3G**). Indeed, these gene expression changes may underlie the observed elevated meiotic recombination rates of placental mammals.

Finally, at post-meiotic stages (three-month opossum and 20 dpp mouse), we identified differences in expression of several genes important for sperm function. *CRISP2*, a gene important for sperm motility and acrosome function in human and mouse (Lim et al. 2019), and *PDE8A*, a gene linked to modulation of Leydig cell function (Vasta et al. 2006), are both more highly expressed in mouse compared to opossum spermatids. In contrast, *DUSP21*, a testis-specific dual-species phosphatase important for the sperm mitochondrial membrane and for microtubules in mature sperm flagella (Hood et al. 2002; Rardin et al. 2008; Leung, Miguel Ricardo et al. 2022), and *TCP11×2*, a gene involved in sperm motility, capacitation, and cAMP signaling (Castaneda et al. 2020; Fraser et al. 1997), are both more strongly expressed in opossum spermatids (**Figure 5C**). The functional significance of these differences for sperm function and divergence in reproductive function remains unexplored.

### Marsupial-placental mammal differences in timing of gene expression

In addition to cases where levels of expression differ quantitatively between species, we also identified multiple genes whose timing of expression during spermatogenesis is different in opossum compared to mouse. Two examples are shown in **Figure 5D**. *FAM71B* shows high expression in opossum spermatocytes but is delayed until the spermatid stage in mouse, while *ROPN1* is expressed in round spermatids in opossum and delayed until the elongating spermatid stage in mouse. *ROPN1* is important for sperm motility and fibrous sheath structure (Fujita et al. 2000; Fiedler et al. 2013).

Divergence in timing along the same developmental trajectory thus represents another modality of gene expression evolution - heterochrony – which is obscured by snapshots of single time-points of fully mature organs but uncovered in a diachronic process like spermatogenesis.

### Gene expression differences identified in testis somatic cells

Many of the genes with elevated expression in opossum compared to mouse (Group 2 in **Figure 3G**) are associated with testis somatic cells rather than germ cells. We also observed a qualitative increase in the proportion of the testis occupied by somatic cells in opossum in both scRNA-seq data (**Figure 1B**) and histological sections (**Figure 1A**, **Figure 4B**). Examination of our adult scRNA-seq data showed that the higher levels of expression of these genes in opossum in bulk RNA-seq data was not a consequence of increased somatic cell abundance, but instead indicated a true difference in gene expression level within the somatic cell populations (**Figure 5E**). For example, INSL3 is a hormone produced by Leydig cells that has roles in the transabdominal phase of testicular descent in early stages of development, puberty and germ cell survival in humans, and is linked to reproductive health and aging (Nef and Parada 1999; Esteban-Lopez and Agoulnik 2020; Ivell et al. 2022, 2013; Zimmermann et al. 1999; Kawamura et al. 2004; Ferlin and Foresta 2005). The *INSL3* gene showed significantly higher expression in opossum compared to mouse somatic cells in most stages of our age-matched comparison, potentially indicating differential function (**Supplemental Table 3**). *GSTM3*, a glutathione S-transferase important for mitigating oxidative stress, promoting germ cell viability, and enabling sperm-oocyte binding (Llavanera et al. 2020), showed significantly higher expression in opossum compared to mouse Leydig and Sertoli cells. Interestingly, several genes involved in controlling oxidative stress were expressed at higher levels in opossum compared to mouse in both somatic and germ cells, and in germ cells this class of genes follows a typical pattern for Group 2 genes with dynamic expression levels across placental species (**Figure 5F**). Oxidative stress pathways are activated in premature infants due to early exposure to an oxygen-rich environment (Cannavò et al. 2021; Frank 1985; O’Donovan and Fernandes 2004), suggesting that relatively high levels of expression of oxidative stress pathways in opossum might help to protect underdeveloped neonates from exposure to reactive oxygen species.

## Discussion

Here, we apply a comprehensive analysis of spermatogenic gene expression in the model marsupial *Monodelphis domestica* (grey short-tailed opossum) and compare it to equivalent data in the mouse and find hundreds of genes whose expression differs between species. We classify these genes into three groups with distinct evolutionary trajectories in spermatogenic gene expression (**Figure 6**). First, we find a class of genes that are expressed in the strongly conserved cell types and developmental events associated with spermatogenesis in amniotes, which include most of the canonical spermatogenesis genes such as *HORMAD2* and *SYCP1*. A second class of genes, including regulators of oxidative stress such as *GSTM3* and *PTGS2*, shows gains and losses of expression along multiple branches of the mammalian clade and may demonstrate species-specific adaptations, providing a substrate for evolutionary experimentation (Kaessmann 2010). Because spermatogenic cells undergo multiple drastic chromatin changes and globally relax transcriptional repression, spermatogenesis represents an opportunity for changes in gene expression to arise and be perpetuated in future generations (Assis and Bachtrog 2013; Findlay et al. 2009; Kaessmann 2010; Kondo et al. 2017; Marques et al. 2005; Xia et al. 2020). Finally, a third class of genes, including regulators of meiotic recombination and DNA repair, gained expression in the placental common ancestor and has maintained elevated expression in spermatogenesis throughout the placental lineage. These genes likely represent placental mammal innovations in gene expression during germ cell development that may contribute to species differences in germ cell function and species-specific developmental traits. Together, the patterns we define here demonstrate how the unusual features of spermatogenic gene expression play out at evolutionary time scales and may be used to elucidate mechanisms of gene regulatory evolution.

**Figure 6:**
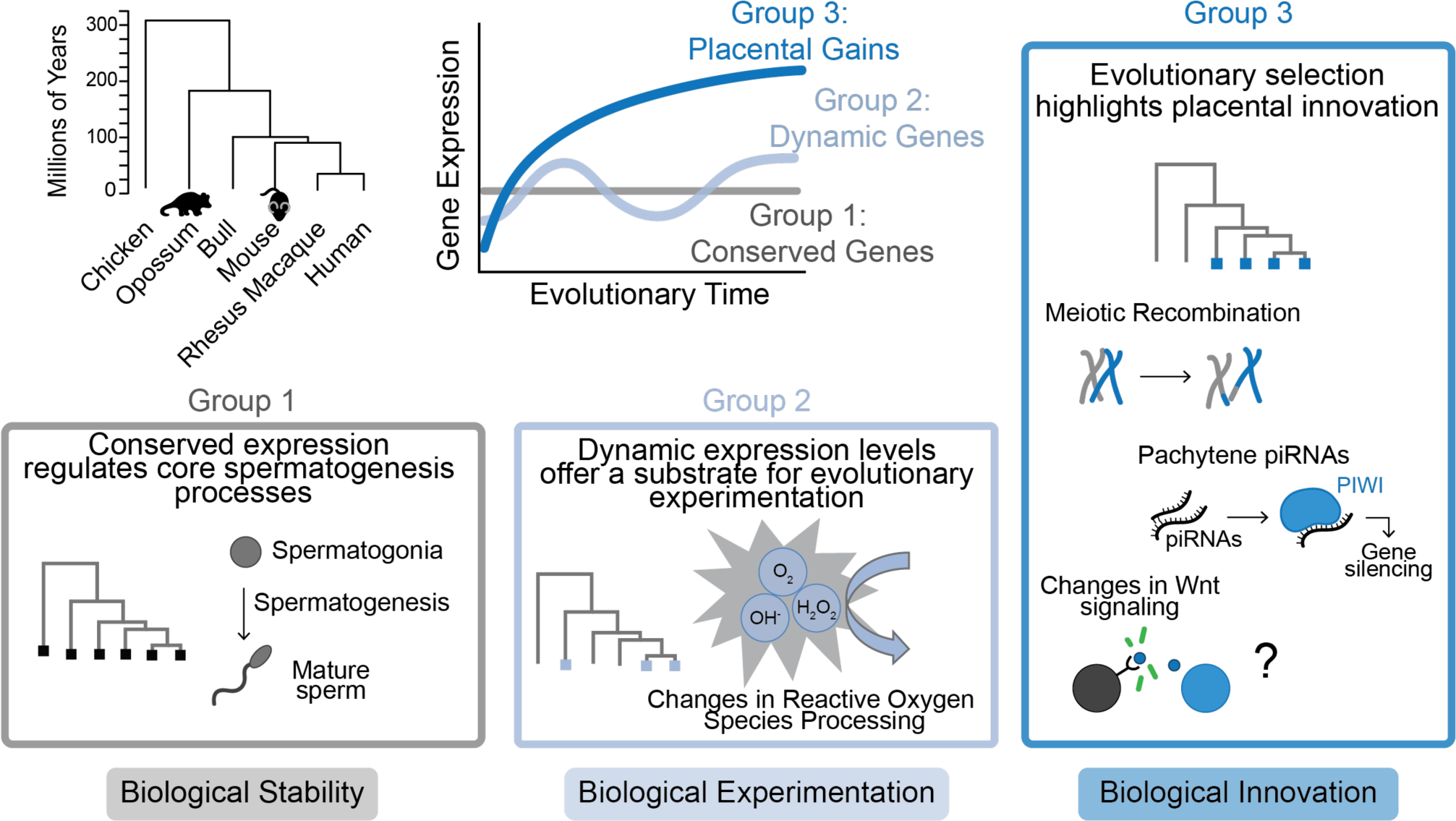
Evolutionary dynamics of gene regulation during spermatogenesis. Summary of evolutionary dynamics of gene regulation during spermatogenesis across the placental-marsupial split, showing three categories of genes revealed by this study: conserved expression (Group 1: black), genes with dynamic expression across mammalian evolution (Group 2: light blue), and directional selection for evolutionary innovations in spermatogenic gene expression in the placental mammal ancestor (Group 3: dark blue). Boxed regions highlight pathways included in each regulatory group.

While the spermatogenic developmental program is strongly conserved across species as expected, we defined numerous specific transcriptional differences at both the molecular and cellular levels during spermatogenesis, pointing to the molecular underpinnings of biological differences between species. An established difference in mouse and opossum germ cell biology is meiotic recombination rate: *Monodelphis domestica* has a much lower rate of recombination events than eutherian mammals (Mikkelsen et al. 2007; Samollow 2008; Samollow et al. 2007). Accordingly, our study identified multiple differentially expressed genes relating to recombination and DNA damage repair, most of which followed a general pattern of higher expression in mouse compared to opossum germ cells and corresponded to our more general Group 3 pattern, with elevated expression across the placental lineage. An example is the kinase ATR, which contributes to synapsis and recombination of homologous chromosomes in mouse (Pacheco et al. 2018; Widger et al. 2018), and we identified as having significantly lower expression in opossum germ cells compared to placental mammals (**Figure 5B, Supplemental Table 3, Supplemental Figure 6F**). Many of these genes were differentially expressed specifically upon addition of meiotic cells during the first wave (at 2 months in opossum and 14 dpp in mouse), emphasizing the developmental specificity of these differences and revealing possible molecular mechanisms.

A second well-characterized difference in mouse and opossum germ cells is the presence of pachytene piRNAs in mouse, a class of regulatory small RNAs unique to germ cells of placental mammals (Chirn et al. 2015; Girard et al. 2006; Grivna et al. 2006; Özata et al. 2020; Sun et al. 2022; Yu et al. 2021). Expression of these piRNAs starts during the pachytene stage of meiosis I in mice and humans, but evidence of their expression has not been found in marsupials, including *M. domestica* (Aravin et al. 2006; Chirn et al. 2015; Girard et al. 2006; Grivna et al. 2006; Özata et al. 2020; Sun et al. 2022; Yu et al. 2021). Correspondingly, we detected lower expression of Argonaute gene *PIWIL2* in opossum and confirmed this finding with RT-qPCR and RNA FISH (**Figure 4**). PIWIL2 is a key protein in piRNA processing, including but not limited to pachytene piRNAs (Grivna et al. 2006; Kuramochi-Miyagawa et al. 2004; Lee et al. 2006). The requirement for PIWIL2 in processing other piRNA classes may explain the observation that it is still expressed in opossum, albeit at reduced levels.

We also identified multiple expression changes likely to represent previously undescribed differences in germ cell molecular function. Genes controlling sperm morphology and sperm-egg interactions are upregulated after meiosis and represent frequent targets of positive selection (Dorus et al. 2010; Swanson and Vacquier 2002; Vicens et al. 2014). Mature opossum sperm pair at the acrosome and swim in tandem, while mouse sperm do not, representing a clear biological difference in function at late spermatogenic stages (Moore and Taggart 1995; Hwang et al. 2021). We found a significant increase in *HEXB* transcript in opossum testes (**Figure 4**), encoding a lysosomal enzyme that is crucial for epidydimal function in mouse (Adamali et al. 1999b, 1999a), suggesting an earlier or divergent role for *HEXB* in male gamete development in opossum. *TCP11×2*, a transcript present in round spermatids and also expressed more strongly in opossum compared to mouse, has been linked to sperm motility, capacitation, and cAMP signaling (**Figure 5C**) (Castaneda et al. 2020; Fraser et al. 1997). Meanwhile, at earlier stages of germ cell development, ubiquitin ligase NEDD4 and RNA binding protein LIN28A both regulate homeostasis of spermatogonial stem cells in mouse and human (Chakraborty et al. 2014; Wang et al. 2020; Zheng et al. 2009; Zhou et al. 2017), and each of these genes shows lower expression in opossum germ cells, with levels of *LIN28A* being almost undetectable (**Supplemental Figure 6F**). We also detected multiple differences in expression of genes participating in the WNT signaling pathway (**Figure 5A**). Such differences could underlie divergence in regulation of spermatogenic stem cells in the testis between marsupials and placentals, an intriguing possibility that will require further investigation. Finally, we found differences in both the quantity and expression levels of somatic cells in the testis, suggesting the presence of previously undescribed differences in molecular function of these somatic cells.

By combining analysis of scRNA-seq and developmentally staged bulk RNA-seq data, we developed a sensitive and robust pipeline for identifying quantitative differences in gene expression within a given cell type. There are several tools available for optimizing scRNA-seq pipelines and clustering analysis across species (Dhodapkar 2022; Li et al. 2022; Pan et al. 2022; Squair et al. 2021; Tarashansky et al. 2021), but no robust tool for quantitative comparison of single cell gene expression levels, especially in cross-species analysis. Our combined approach enabled us to identify novel differences in gene expression levels between mouse and opossum spermatogenic cells with high confidence. We observed a tendency toward detection of higher expression in mouse compared to opossum among differentially expressed genes; this effect may be a reflection of the differing evolutionary dynamics identified between Group 2 and Group 3 genes (**Figure 6**), or it may be due to differences in genome annotation quality between species. The possibility of annotation-related artifacts emphasizes the importance of experimental validation for differential expression calls and suggests that our analysis may miss some cases where a gene is more strongly expressed in opossum compared to mouse. Our validation of selected genes by RT-qPCR and RNAScope in this study argues that despite imperfect annotation, many of our differential gene calls reflects true *in vivo* differences in expression. This conclusion is further supported by the correspondence between differential expression and differential enrichment of the activation-associated modification H3K4me3 (**Figure 3D-F**): globally, genes with higher expression in one species also had higher H3K4me3 enrichment at their promoters. Our data thus suggest that differences in gene expression can be used as a proxy for identifying divergence in underlying chromatin states and is a useful starting point in investigating divergence in regulatory mechanisms across species.

Together, our study leverages histology, scRNA-seq, and bulk RNA-seq data to define conserved and divergent gene expression programs across the mammalian lineage. We define the timing of first-wave spermatogenesis in opossum and provide a robust comparison to spermatogenesis in mouse. We establish the coexistence of contrasting forces underlying the evolutionary dynamics of spermatogenic gene expression by identifying three classes of genes with different evolutionary trajectories: a deeply conserved central gene regulatory program governing spermatogenic progression; a set of genes whose expression levels during spermatogenesis fluctuate across placentals indicating dynamic species adaptations; and a set of genes with evidence for directional selection at the base of the placental mammalian lineage that represent placental innovations in germline gene expression. Our data highlight potential biological innovations in germ cell development, provide a resource for fueling future research into functional species differences in germ cell biology and developmental phenotypes, and create a framework for studying cross-species transcriptomics and regulatory evolution more generally.

## Methods

### Animal care

These studies were approved by the Yale University Institutional Animal Care and Use Committee under protocol 2020-20169 and 2021-11483. Gray short-tailed opossums (*Monodelphis domestica*) were raised in a breeding colony according to established protocols (Keyte and Smith 2008) and ethical guidelines approved by the Yale University Institutional Animal Care and Use Committee. Mice were kept under standard conditions and all experiments were conducted in compliance with the Animal Welfare Act. All animals used in these studies were maintained and euthanized according to the principles and procedures described in the National Institutes of Health Guide for the Care and Use of Laboratory Animals.

### Sample acquisition and processing

#### Whole testis dissociation

Testes from gray short-tailed opossums at Yale or adult (8-16 week old) male C57BL6/J mice purchased from Jackson Laboratories were harvested and dissociated into single-cell suspension for RNA isolation using collagenase and trypsin. Briefly, testes were removed from the scrotum, decapsulated from the tunica albuginea, and incubated in 0.75mg/mL collagenase in Dulbecco’s Modified Eagle Medium (Gibco 11965-09L; Thermo Fisher Scientific, Inc.) at 37°C. DMEM was added to dilute collagenase and samples were centrifuged for 5 minutes at 400 x g at room temperature. Supernatant was discarded and samples were washed in DMEM. Samples were resuspended in 0.05% trypsin-EDTA (Gibco 25300-054; Thermo Fisher Scientific Inc.) with DNAse (1:10,000; Stem Cell Technologies #07900, Vancouver BC, Canada). The reaction was quenched with 10% Cosmic Calf Serum (Sigma-Aldrich #C8056, Burlington, MA, USA) in DMEM (CCS-DMEM) and centrifuged. Samples were resuspended in CCS-DMEM and filtered through a 100-micron filter. Cell concentration was determined by hemocytometer.

#### RNA isolation

Aliquots of whole testis single-cell suspension were centrifuged at 6000 x g for 4 minutes at 4°C. Samples were resuspended in 500 μL Trizol (Ambion #15596018, Thermo Fisher Scientific, Inc.) and disrupted by drawing up and down through at 21-gauge needle and syringe. The aqueous layer was separated using 50 μL 1-bromo-3-chloropropane (BCP; Sigma-Aldrich #B9673, Burlington, MA, USA) per 500 μL of Trizol and centrifuged at 12000 x g for 15 minutes at 4°C. Genomic DNA was removed using the genomic DNA eliminator columns supplied in the Qiagen RNeasy Plus Mini kit (74134, Hilden, Germany). All remaining steps of RNA isolation were performed according to the manufacturer’s instructions using the RNeasy Plus Mini kit. Samples were processed in batches of 1-5 in order of collection and stored at −80°C.

### Bulk RNA-seq

Bulk RNA-seq libraries were prepared using either KAPA mRNA HyperPrep Kit (Roche # 08098123702) or KAPA RNA HyperPrep Kit with RiboErase (HMR) (Roche # 08098140702) and sequenced on an Illumina NovaSeq to generate paired-end, 100 base pair read libraries at an average depth of 35 million read pairs/library. Bulk RNA-seq data was generated from dissociated whole testes from opossum collected at one month (n = 2), two months (n = 4), three months (n = 4), four months (n = 3), or adult (i.e. > 6 months; n = 7). Mouse RNA-seq data was obtained from Margolin et al. (Margolin et al. 2014) (GEO accession GSE44346). These data were 36 base pair, single-end reads from testes of animals at 6 days postpartum (dpp; n = 6), 10 dpp (n = 5), 12 dpp (n = 6), 14 dpp (n = 6), 16 dpp (n = 13), 18 dpp (n = 6), 20 dpp (n = 5), or adult (i.e., 38 dpp; n = 8).

### Bulk RNA-seq analysis

RNA-seq libraries were aligned using HISAT 2.2.1(Kim et al. 2019) with Ensembl(Yates et al. 2020) release 104 as a reference and data output type. Output files were converted from SAM format to sorted BAM format using SAMtools-1.12 (Danecek et al. 2021; Li et al. 2009). Raw count values and FPKM were obtained using StringTie 2.1.4 (Pertea et al. 2016) with the -e and -G options and Ensembl (Yates et al. 2020) ASM229 release 104 or GRCM39 release 104 reference files for opossum and mouse, respectively.

### Single cell RNA-seq

Whole testis single cell suspension from opossum or mouse (n = 2 for each species) was diluted to 10,000 cells/mL and single cell RNA-seq libraries were prepared using the Chromium Next GEM Single Cell 3’ GEM Library Kit v. 3.1 (10x Genomics #PN-1000121). Libraries were sequenced on an Illumina NovaSeq to generate paired-end, 75 or 150 base pair read libraries at an average depth of 250 million read pairs/library.

### Single cell RNA-seq analysis

Single cell RNA-seq library alignment to Ensembl GRCm39 (mouse) or ASM229v1 (opossum) and quality control was performed using 10x Genomics Cell Ranger 6.0.1 with default parameters (Zheng et al. 2017). CellRanger filtered matrices were then used for further filtering with SoupX (Young and Behjati 2020) using the standard settings and no additional clustering data. Strained SoupX count matrices were used for all downstream analyses.

#### Filtering, doublet removal, and integration of replicates

Filtered and strained count data for mouse and opossum were analyzed with Scanpy 1.9.1 (Wolf et al. 2018). Starting datasets included 10,586 cells and 12,801 for mouse replicates and 8,745 and 9,602 for opossum replicates. Due to species-specific restrictions with calling mitochondrial gene percentage we chose to filter for high quality cells using the number of unique genes per cell. To remove likely empty or contaminated droplets, we filtered out any cells below the 15^th^ percentile or above the 98^th^ percentile of number of genes expressed (**Supplemental Figure 1B**). We then used DoubletDetection v4.2(Gayoso and Shor 2019) to remove any remaining doublets. Each dataset was then normalized using the standard log(x+1) and a further normalized data layer for gene expression visualization was added using square-root transformation followed by Markov Affinity-based Graph Imputation of Cells (MAGIC; (van Dijk et al. 2018)). After filtering, each dataset contained 8,282 or 10,502 cells (mouse) and 7,160 or 7,663 cells (opossum). The two replicates were then combined within each species using Harmony v1.0 (**Supplemental Figure 1C-D**, (Korsunsky et al. 2019). All further analyses were performed using the harmony-combined data. Final combined datasets contained 18,784 cells or 14,823 cells for mouse and opossum, respectively.

#### Cell clustering and selection of germ cells

Datasets were embedded using the Scanpy neighbors function with ten neighbors and clustered using the leiden function (**Supplemental Figure 1E,** (Virshup et al. 2023)). Previously established cell-type marker genes in mouse were used to broadly identify cell cluster identity (**Supplemental Table 1**, **Supplemental Figure 1E, 2-4,** (Hermann et al. 2018; Green et al. 2018; Shami et al. 2020; Wang et al. 2018; Murat et al. 2023)). Further analysis was performed using only clusters representing germ cells (9,548 cells and 4,548 cells for mouse and opossum, respectively) or somatic cells (1,405 cells and 4,249 cells for mouse and opossum, respectively). Subsetted cell data was re-embedded and clustered using the leiden function (**Supplemental Figure 1E**). Clusters were re-assigned cell-type identity using cell-type marker genes (**Supplemental Table 1, Supplemental Figure 3, 4)**. The germ cell cluster labeled ‘transitional’ was removed from all further analyses, filtering the datasets to 9,112 cells and 2,990 cells for mouse and opossum, respectively. Cluster designations were compared between species using SAMap (**Supplemental Table 1,** (Tarashansky et al. 2021)). For each cluster, the top 100 marker genes were determined using the Wilcoxon rank-sum function. Separately, we used Potential of Heat-diffusion for Affinity-based Trajectory Embedding (PHATE; (Moon et al. 2019)). with settings k=5, a=20, t=150 as an alternative method to embed data while preserving progression structure, confirming marker progression observed in the previous analysis. Gene expression maps were scaled for each gene with “low” being zero and “high” being maximum expression value.

### Validation of cell clustering with sorted germ cell data

Gene expression data for sorted pachytene spermatocyte and round spermatid cells from mouse and opossum were obtained from Lesch et al. (Lesch et al. 2016); GEO accession GSE68507)). FPKM values were converted to zFPKM values according to Hart et al (Hart et al. 2013) using log_2_ transformed values of FPKM + 0. 00001 to avoid removal of genes with no expression in only one species:

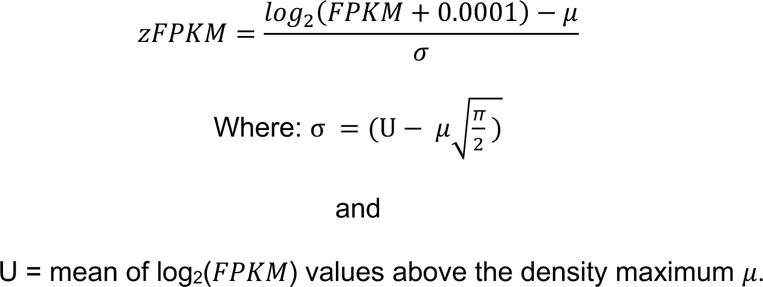

Following normalization, genes identified as markers of spermatocyte or round spermatid clusters from single cell analysis were selected and normalized expression values for these gene sets were compared between sorted cell populations. Statistical comparisons were performed using a Mann-Whitney U Test with α < 0.05 to indicate significance.

### Histological sections

Whole testes from opossum were fixed in Hartman’s Fixative (Sigma-Aldrich #H0290, Burlington, MA, USA) overnight at 4°C, then transferred to 70% Ethanol. Testes were cross-sectioned followed by dehydration and paraffin embedding. Paraffin sections were mounted on slides and stained with hematoxylin and eosin to visualize DNA and cell structure. Scans of whole testis sections were taken at 40x magnification using a Leica DM6B with attached camera Leica DC310fx in brightfield with brightness settings standardized across all images.

### Differential expression analysis across developmental ages in opossum

Differentially expressed genes between developmental stages of opossum testes were determined via DESeq2 (Love et al. 2014). A raw count matrix containing all ages was input with age as the condition. Principal component analysis was performed using vst-normalized data. Each pairwise comparison was performed by extracting two age groups at a time as contrast. Within each pairwise analysis, p-value was adjusted using false discovery rate. Data was considered significant if the adjusted p-value was below 0.05 and the log_2_FoldChange was greater than 2.0. Differentially expressed genes were extracted and further filtered based on expression to remove inflation caused by low expression levels by retaining only genes with an average FPKM of 50 or higher across all individuals of both ages (**Supplemental Figure 5C**). This threshold was chosen based on FPKM values of genes previously demonstrated to be expressed or functionally off in testis tissues (**Supplemental Figure 5D**).

### Comparison of developmental differentially expressed genes to scRNA-seq clusters

Genes identified as differentially expressed with the addition of spermatocytes (1 month vs 2 month) or round spermatids (2 months vs 3 months) were selected and their expression was compared to average cluster expression or cluster markers from adult scRNA-seq. For average cluster expression, a pseudobulk profile was created using the get_pseudobulk function of decoupleR (Badia-I-Mompel et al. 2022) with mode = sum and gene expression averages for genes upregulated at two months or three months in the DESeq2 analysis were extracted for every cluster. Statistical comparisons were performed using a Kruskal-Wallis rank sum test followed by a Dunn test for multiple comparisons with Bonferroni correction.

For comparison to cluster markers, the top 100 cluster markers for spermatocyte, late spermatocyte, round spermatid, and late round spermatid clusters were determined using the Wilcoxon rank-sum function (**Supplemental Table 1**). Cluster marker gene lists were reduced to only include genes also represented in the DESeq2 analysis. The percentage of cluster marker genes identified as differentially expressed was calculated for spermatocyte or late spermatocyte clusters with genes upregulated at 2 months and for round spermatid or late round spermatid clusters with genes upregulated at 3 months. To determine similarities between developmental differentially expressed genes and somatic cell clusters, the top 100 cluster markers for each somatic cell type cluster were identified and combined into a final list of 594 marker genes (**Supplemental Table 2**). The percentage of cluster marker genes identified as differentially expressed was calculated for with genes upregulated at 2 months or 3 months separately.

### Comparison of gene expression across species

Only genes with one-to-one orthologs between mouse and opossum that aligned to a known chromosome were included. One-to-one orthologs were obtained using the BioMart(Durinck et al. 2005, 2009) database with Ensembl release 104 (Yates et al. 2020). Gene lists were filtered to remove genes with FPKM of zero in all samples (both species). A final set of 13,521 genes was used for cross-species analysis (**Supplemental Table 3**).

#### Pst calculation

*P_st_* values are a comparative measure of variation within between species for a quantitative trait such as gene expression (Antoniazza et al. 2010; Uebbing et al. 2016). Calculations were performed as described in Uebbing et al. (Uebbing et al. 2016) after determining zFPKM to represent expression values (**Supplemental Table 5**) with an assumed h^2^ value of 1, corresponding to the value used previously in similar analyses (Antoniazza et al. 2010; Uebbing et al. 2016). P values were obtained by bootstrapping for 1000 iterations to determine likelihood of obtaining the observed value and then adjusted for multiple comparisons using the Benjamini-Hochberg procedure. To determine high-confidence differentially expressed genes, we added a stringent filter for between-species variation values to minimize false positives. We set between-species variation thresholds by evaluating previously characterized genes known to have either similar or significantly different expression, including stably expressed genes elongation factor-2 (*EEF2*) and β-actin (*ACTB*) and differentially expressed genes histone methyltransferase *PRDM9* and its cofactor *ZCWPW1* (Baker et al. 2017; Cavassim et al. 2022; Sun et al. 2015) (**Supplemental Figure 6A-B**). Genes with an adjusted p-value less than 0.05 and a between-species variation greater than 3.5 were considered differentially expressed between mouse and opossum.

#### Determining age-matched timepoints for cross-species comparisons

We used our histology data to inform possible matching opossum and mouse time points based on when a given cell type is first observed (**Figure 2A**) (Bellvé et al. 1977; McCarrey 1993). For each opossum age, we performed two pairwise Pst comparisons at the two best-matched mouse ages based on histology, with the exception of the one-month time point in opossum that only had one possible equivalent (6dpp) in the mouse first-wave dataset. We determined the best time equivalents as the pair with the lower number of differentially expressed genes called between the two possible mouse comparisons and used the resulting paired time points for further analysis (**Figure 3B, 3C**; **Supplemental Table 3**).

### Comparison of developmental differentially expressed genes to chromatin data in sorted germ cells

H3K4me3 signal from chromatin immunoprecipitation followed by sequencing (ChIP-seq) in promoter regions for STA-PUT sorted pachytene spermatocyte and round spermatid cells from mouse and opossum were obtained from Lesch et al. (Lesch et al. 2016) (GEO accession GSE68507) and normalized as described above. Genes identified as differentially expressed at any age group between opossum and mouse were selected and the enrichment of H3K4me3 was compared between species in the same cell types. Statistical comparisons were performed using a Mann-Whitney U Test with α < 0.05 to indicate significance. Metagene analysis was performed using deepTools 3.5.1 computeMatrix function with reference point TSS and 2000 bases each direction (Ramírez et al. 2016).

### Identifying evolutionary innovations in placental mammalian gene expression

Gene expression data for sorted pachytene spermatocyte and round spermatid cells from human (*Homo sapiens)*, rhesus macaque (*Macaca mulatta*), mouse (*Mus musculus*), bull (*Bos taurus*), opossum (*Monodelphis domestica*), and chicken (*Gallus gallus*) were obtained from Lesch et al. (Lesch et al. 2016) (GEO accession GSE68507). FPKM values were converted to zFPKM values as described above. Following normalization, genes identified by cross-species Pst analysis as showing conserved expression, higher expression in mouse, or higher expression in opossum were selected and zFPKM values for these gene sets were compared between sorted cell populations. PhastCons and PhyloP conservation scores were calculated with reference point TSS and 1000 bases each direction using bigWigAverageOverBed function and Vertebrate Multiz Alignment and Conservation of 35 vertebrate genomes to the mouse genome (phastCons 35way or phyloP 35way) (Yates et al. 2020; Kent et al. 2010; Blanchette et al. 2004; Siepel et al. 2005). Statistical comparisons were performed using a Kruskal-Wallis rank sum test followed by a Dunn test for multiple comparisons with Bonferroni correction.

### Reverse transcription and quantitative PCR

Reverse transcription of RNA isolated from dissociated whole testis of mouse and opossum at each developmental time point was performed with oligo dT primers and SuperScript III reverse transcriptase (Invitrogen #18080-051, Thermo Fisher Scientific, Inc.) according to the manufacturer’s instructions. Reaction mixtures (20 μL) were incubated in a thermocycler for 50 minutes at 50°C followed by 5 minutes at 85°C to stop the reaction. cDNA was then diluted 1:5 in nuclease-free water. Real-time quantitative PCR reactions were performed in 20 μL reactions consisting of 4 μL diluted cDNA, 10 μL Power SYBR Green PCR Master Mix (Applied Biosystems 4367659, Thermo Fisher Scientific, Inc.), 5.6 μL nuclease-free water, and 0.4 μL of 10 μM forward and reverse primer mixture (**Table 1**). Reactions for each target were performed in duplicate in a 96 well plate using a QuantStudio 3 Real-Time PCR System (Applied Biosystems, Thermo Fisher Scientific, Inc.) with standard cycling conditions (Hold stage (x1): 50°C for 2 min, 95°C for 10 min; PCR stage (x40): 95°C for 15 sec, 60 °C for 1 min, Melt curve stage: 95°C for 15 secs, 60°C for 1 min, 95°C for 15 secs). Relative expression was calculated as 2-^ΔΔCt^ where ΔC_t_ = mean of target C_t_ values – mean of control *EEF2* C_t_ values within each species and ΔΔCt = ΔC_t_ – [mean(ΔC_t_) of both species combined]. Significance was calculated by unpaired t-test.

**Table 1:**
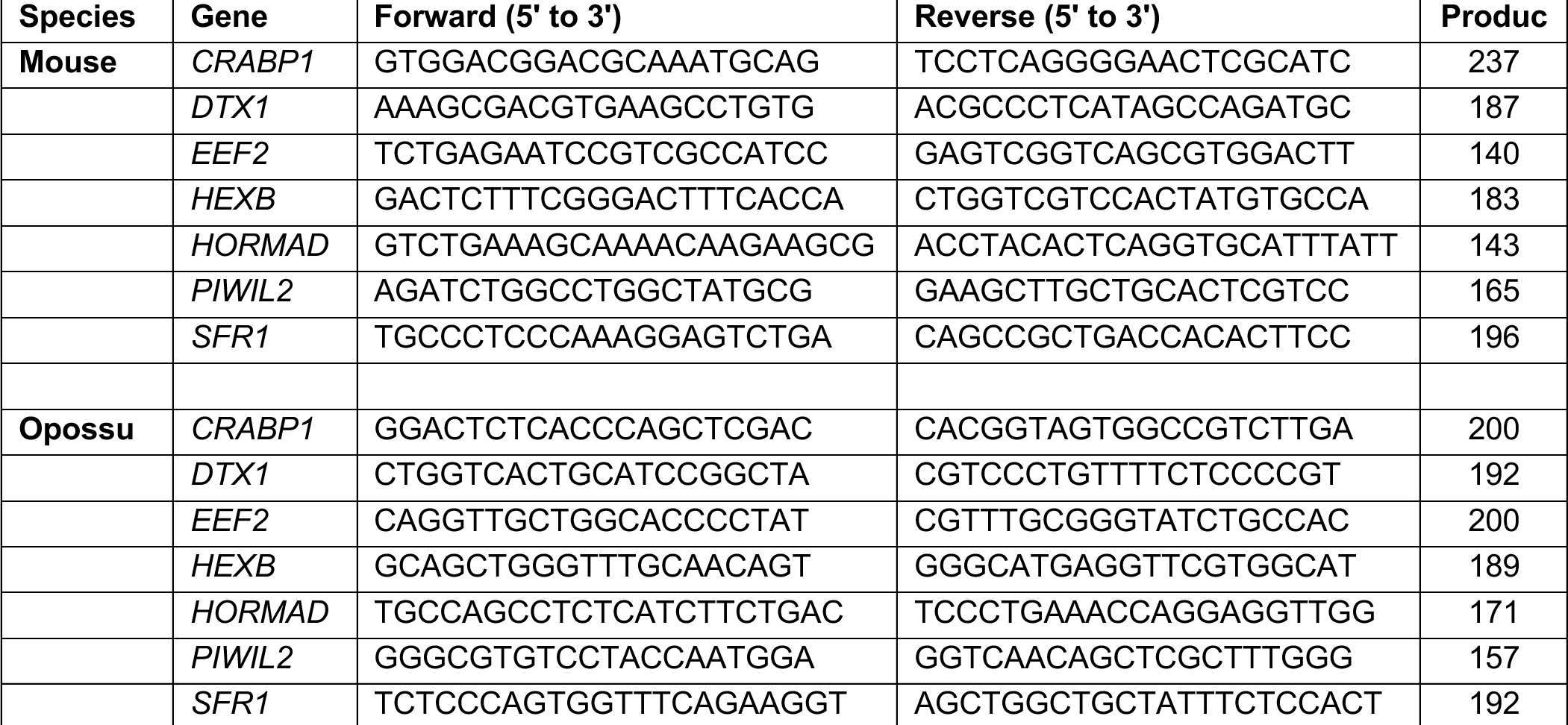
qPCR Targets and Primers.

### RNAscope

Whole testes from adult mouse and opossum were harvested and fixed in 4% paraformaldehyde overnight at 4°C. Samples were dehydrated through an ethanol gradient followed by two incubations in xylenes. Paraffin infiltration in three 20-minute incubations was performed followed by paraffin embedding. Blocks were allowed to solidify at room temperature prior to sectioning using a microtome and mounting on glass slides. Slides were dried overnight at room temperature prior to beginning RNAscope. Slides were then stained using RNAscope 2.5 HD Detection Kit-RED (#322350, ACD Bio, Newark, CA) per the manufacturer’s instructions. Sample probes targeting *HEXB* and *PIWIL2* as well as a positive control collagen gene *COL3A1* and a negative control targeting bacterial gene *dapB* were used (**Table 2**; ACD Bio, Newark, CA), with probes designed separately for the mouse and opossum orthologs of each gene. Scans of whole testis sections were taken at 40x magnification using a Leica DM6B with attached camera Leica DC310fx in brightfield with brightness settings standardized across all images.

**Table 2:** RNAscope Targets and Probes.

| Species | Gene | Probe ID |
| --- | --- | --- |
| Mouse | *HEXB* | 314231 |
|  | *PIWIL2* | 177541-C1 |
|  | *COL3A1* (Positive) | 455771 |
| Opossum | *HEXB* | 1177521-C1 |
|  | *PIWIL2* | 1177511-C1 |
|  | *COL3A1* (Positive) | 1147071-C1 |
| Negative | *dapB* | 310043 |

## Data Access

All raw and processed RNA-seq data generated in this study have been submitted to the NBCI Gene Expression Omnibus (GEO; https://www.ncbi.nlm.nih.gov/geo/) under accession number GSE216343. Code used for analysis is available on Github (https://github.com/Lesch-Lab).

## Acknowledgments

We would like to thank Dr. Severin Uebbing for his assistance with zFPKM normalization and Cody Limber and Dr. Kim Griffin for help with image acquisition. We thank Shannon Rainsford and Dr. Zachary Smith for opossum scRNA-seq library preparation, and Dr. Aushaq Malla for help evaluating testis histology. We thank Delaney Farris for her help with graphical designs. We also thank the Yale Comparative Pathology Research Core for histological sample preparation and the Yale Center for Genomic Analysis for RNA sequencing services.

This work was supported by funding from the National Institute of Child Health and Human Development (NICHD, R01HD098128), the Kinship Foundation through the Searle Scholars Program, and a Pew Biomedical Scholar Award from the Pew Charitable Trusts to B.J.L. K.L.M is supported by an NIH predoctoral fellowship from NICHD under award number F31HD107950. K.L.M and D.J.S received funding from the NIGMS Genetics Predoctoral Training Program (T32 GM7499-43). Research in the G.P.W lab is supported by John Templeton Foundation grant ID#61329 and Burroughs Wellcome Fund ID#1022864; this article reflects the opinions of the authors and not that of the John Templeton Foundation.

## Author contributions

Conceptualization: K.L.M., B.J.L.

Software: D.J.S., K.L.M.

Validation: K.L.M.

Formal Analysis: K.L.M.

Investigation: J.M., D.J.S., K.L.M.

Resources: G.P.W., B.J.L.

Writing – Original Draft: K.L.M.

Writing – Review & Editing: D.J.S., G.P.W., B.J.L.

Visualization: K.L.M.

Supervision: B.J.L.

Funding acquisition: G.P.W., B.J.L.

